# Perturbation-response genes reveal signaling footprints in cancer gene expression

**DOI:** 10.1101/065672

**Authors:** Michael Schubert, Bertram Klinger, Martina Klünemann, Mathew J Garnett, Nils Blüthgen, Julio Saez-Rodriguez

## Abstract

Aberrant cell signaling is known to cause cancer and many other diseases, as well as a focus of treatment. A common approach is to infer its activity on the level of pathways using gene expression. However, mapping gene expression to pathway components disregards the effect of post-translational modifications, and downstream signatures represent very specific experimental conditions. Here we present PROGENy, a method that overcomes both limitations by leveraging a large compendium of publicly available perturbation experiments to yield a common core of Pathway RespOnsive GENes. Unlike existing methods, PROGENy can (i) recover the effect of known driver mutations, (ii) provide or improve strong markers for drug indications, and (iii) distinguish between oncogenic and tumor suppressor pathways for patient survival. Collectively, these results show that PROGENy more accurately infers pathway activity from gene expression than other methods.

## Introduction

A wealth of molecular data has become available to describe a cell’s state in different diseases. The remaining challenge is how to derive predictive and reliable biomarkers for disease status, treatment opportunities, or patient outcome in a way that is both relevant and interpretable. Of particular interest are methods which infer and quantify deregulation of signaling pathways, as those are key for many processes underpinning different diseases. Here we focus on cancer, which is largely caused by cell signaling aberrations created by driver mutations and copy number alterations (Hanahan and Weinberg 2000).

Efforts like the TCGA (The Cancer Genome Atlas Research Network et al. 2013) and ICGC have pioneered molecular characterization of primary tumors on a large scale. The GDSC (Garnett et al. 2012; Iorio et (International Cancer Genome Consortium et al. 2010) and CCLE (Barretina et al. 2012) have focussed on preclinical biomarkers of drug sensitivity in cancer cell lines. These initiatives have provided profound insight in the molecular markup of the disease. However, putting the genomic alterations investigated in the functional context of the pathways they alter may shed additional light on mechanisms of pathogenesis and treatment opportunities (Mutation Consequences and Pathway Analysis working group of the International Cancer Genome Consortium 2015).

With direct measurements of signaling activity not widely available, pathway levels have mostly been inferred using the expression of predefined gene sets derived from Gene Set Enrichment Analysis (Subramanian et al. 2005) on Gene Ontology categories (Gene Ontology Consortium 2004) or pathway resources such as Reactome (Croft et al. 2011). More sophisticated methods have attempted to quantify the signal flow by taking into account pathway structure, the best known of which are SPIA Tarca et al. 2008), PARADIGM (Vaske et al. 2010), and Pathifier (Drier et al. 2013). All of these methods however are based on mapping transcript expression on the corresponding signaling proteins, and hence do not take into account the effect of post‐translational modifications that are known to govern mammalian signal transduction (Fig. 1a). It is therefore unclear if and under what circumstances the pathway scores obtained by these methods reflect signaling activity.

**1.**
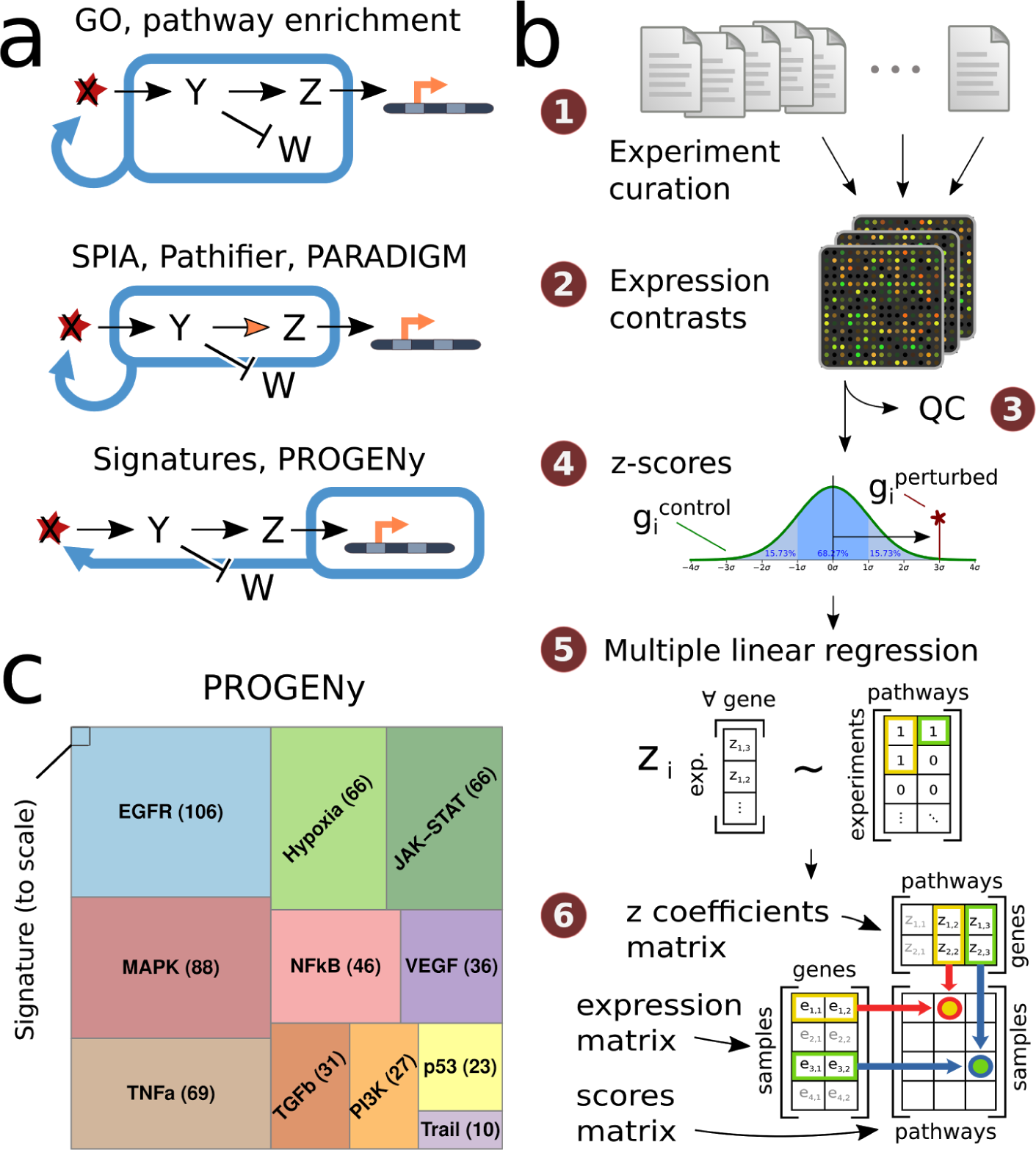
Deriving pathway-response signatures for 11 pathways. A. Reasoning about pathway activation. Most pathway approaches make use of either the set (top panel) or network (middle panel) of signaling molecules to make statements about a possible activation, while our approach considers the genes affected by perturbing the pathway .B. Workflow of data curation and model building. (1) Finding and curation of 208 publicly available experiment series in the ArrayExpress database, (2) Extracting 556 perturbation experiments from series’raw data, (3) Performing QC metrics and discarding failures, (4) Computing z-scores per experiment, (5) Using a multiple linear regression to fit genes responsive to all pathways simultaneously obtaining the z-coefficients matrix, (6) Assigning pathway scores using the coefficients matrix and basal expression data. See methods section for details. Image credit Supplemental Note 1. C. Size of the data set compared to an individual gene expression signature experiment. The amount of experiments that comprise each pathway is shown to scale and indicated.

Another approach is to look at the downstream effect of pathway activity on gene expression. Expression levels of genes regulated by transcription factor or kinases have been used to estimate the activity status of proteins (Chen et al. 2011; Alvarez et al. 2016). Similarly, the transcripts altered when perturbing a specific pathway can be used to infer pathway activation from gene(Bild et al. 2005; Gatza et al.2010), but they are known to be heterogeneous and not replicate well under different experimental conditions (Chibon 2013). This property makes them unsuitable as a generally applicable pathway method.

Here, we overcome the limitations of both approaches by leveraging a large compendium of publicly available perturbation experiments that yield a common core of Pathway RespOnsive GENes to a specified set of stimuli. We then used those to infer the upstream signal mediating downstream expression changes based on building a consensus model (PROGENy), improving on an idea we previously suggested (Parikh et al. 2010).

We performed a systematic comparison of PROGENy and other commonly used pathway methods for 11 cancer-relevant pathways. We investigated how well each method can recover pathway perturbations and is able to recover constitutive activity mediated by driver mutations in The Cancer Genome Atlas (TCGA) (The Cancer Genome Atlas Research Network et al. 2013). We further examined how well they can explain drug sensitivity to 265 drugs in 805 cancer cell lines in the Genomics of Drug Sensitivity in Cancer (GDSC) (Garnett et al. 2012; Iorio et al. 2016) and patient survival in 7254 primary tumors spanning 34 tumor types using TCGA data. We found that PROGENy significantly outperforms existing methods for all these tasks.

## Results

### Consensus gene signatures for pathway activity

We curated (workflow in Fig. 1b) a total of 208 different submissions to ArrayExpress/GEO, spanning perturbations of the 11 pathways EGFR, MAPK, PI3K, VEGF, JAK-STAT, TGFb, TNFa, NFkB, Hypoxia, p53 (and DNA damage response) and Trail (apoptosis). Our dataset consists of 580 experiments and 2652 microarrays, making it the largest study of pathway signatures to date (Fig. 1c and Supplemental Fig. 1).

We obtained z-scores of gene expression changes for each experiment, for which we performed a multiple linear regression using the perturbed pathway as input and gene expression as a response variable. For each pathway, we identified 100 responsive genes that are consistently deregulated across experiments (Supplemental Fig. 2). These responsive genes are specific to the perturbed pathway (Supplemental Fig. 3) and do almost not overlap with genes that comprise it (Supplemental Fig. 4). We use the z-scores of those pathway-responsive genes in a simple, yet effective linear model to infer pathway activity from gene expression called PROGENy (for Pathway RespOnsive GENes, but also to indicate the descent of the method from previously published experiments; Supplemental Table 1).

Had we applied the same methodology on individual signatures at the same significance threshold (10% FDR), those would resemble more the experimental conditions they were derived from than the perturbed pathway (Fig. 2a, left). Instead, applying PROGENy on the input experiments assigns pathway scores that are cluster the input experiments by their intended activation pattern (Fig. 2a, right), suggesting that our model is able to capture the common pathway responses in a heterogeneous set of experiments.

**2.**
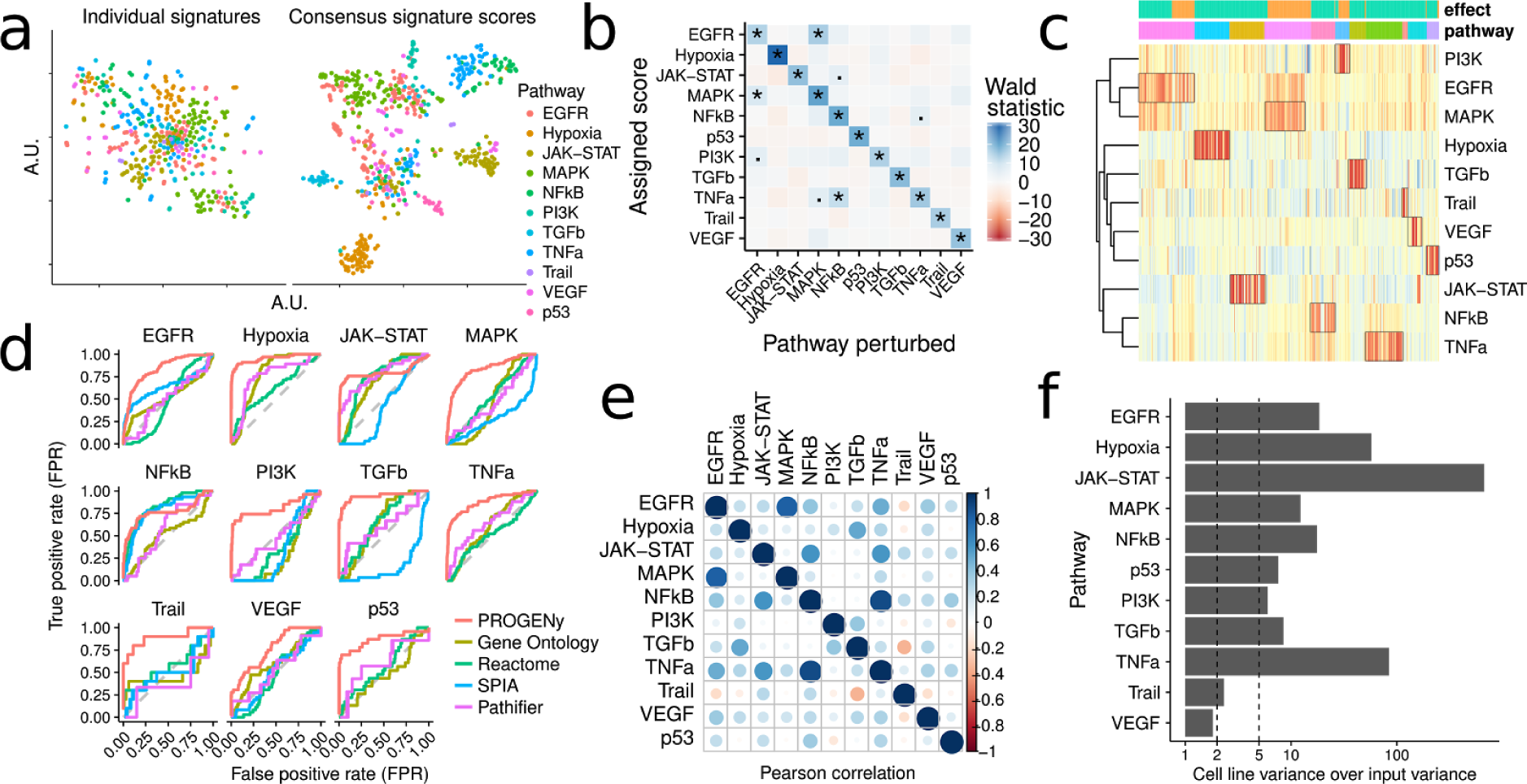
Evaluation of pathway-response signatures. A. T-SNE plots for separation of perturbation experiments with different pathway perturbations in different colors. Fold changes of genes in individual perturbation experiments (10% FDR) do not cluster by pathway (left). Using a consensus signature of genes whose z-score is most consistently deregulated for each pathway instead, we can observe distinct clusters of perturbed pathways (right). Details Supplemental Note 2. B. Associations between perturbed pathways and the scores obtained by the model of pathway-responsive genes (PROGENy). Along the diagonal each pathway is strongly (p<^−10^) associated with its own perturbation. Significant off-diagonal elements are sparse and only occur (p<10^−5^) where there is biologically known cross-activation. C. Heatmap of relative pathway scores in each perturbation experiment. 523 experiments in columns, annotated with the perturbation effect (green for activation, orange for inhibition) and pathway perturbed (same order as b). Pathway scores in rows cluster between EGFR/MAPK and to a lesser extent PI3K, and TNFa/NFkB. Color indicates activation or inhibition strength. D. ROC curves for different methods ranking perturbation experiments by their pathway score. PROGENy show better performance for all pathways except JAK-STAT and NFkB, where other methods are equal. Gene Ontology and Reactome scores obtained by Gene Set Variation Analysis (GSVA). Pathifier using Reactome gene sets. E. Correlation of pathway scores in basal gene expression of cell lines in the GDSC panel. Positive correlation in blue, negative in red. Circle size and shade correspond to correlation strength. Pathways that showed cross-activation in point b are more highly correlated in basal expression as well. F. Stability of basal pathway scores when bootstrapping input experiments. The variance of pathway scores in cell lines given bootstraps more than five times as high compared to the variance of bootstraps given cell lines for all pathways except two (Trail and VEGF), where it is roughly twice as high.

Within experiments, our inferred pathway activation is strongly (p<10^−10^) associated with the pathway that was experimentally perturbed. The associations with other pathways are weaker (p>10^−5^) except for EGFR with MAPK/PI3K and TNFa with NFkB/MAPK (Fig. 2b), where there is biologically known cross-activation (Kant et al. 2011). Relative activation patterns are consistent across input experiments and not driven by outliers (Fig. 2c). This is in contrast to methods based on pathway expression that are not able to recover experimental perturbations by means of their inferred pathway score (Supplemental Fig. 5). Across experiments, PROGENy is able to better rank the perturbations for 10 out of 11 pathways (Fig. 2d and Supplemental Table 6). With the exception of NFkB and JAK-STAT, competing methods do not perform significantly better than random. VEGF is not recapitulated well by any method, possibly because of overlap with other pathways. Overall, PROGENy more closely corresponds to pathway activation upon perturbation than any method that maps transcript expression to pathway members.

Knowing how pathway-responsive genes behave when a stimulus is present, we can take the idea one step further and hypothesize that the existence of a different basal expression level of the responsive genes may in turn correspond to cell-intrinsic signaling activity. We find that the correlation between different pathway scores in basal expression (Fig. 2e) corresponds to the previously observed cross-activation upon perturbation (Fig. 2b and Supplemental Fig. 6), suggesting that PROGENy can detect footprints of signaling activity in basal gene expression. Furthermore, the pathway scores we derive are robust to changes in the experiments that the model was derived from (Fig. 2f).

### Recovering mechanisms of known driver mutations

If our reasoning is correct and pathway-response signatures in basal gene expression correspond to intrinsic signaling activity, we should be able to see a higher pathway score in cancer patients with an activating driver mutation in that pathway and a lower score for pathway suppression compared to patients where no such alteration is present.

We selected all cancer types in the TCGA for which there were tissue-matched normals available, in order to make full use of the pathway methods that require them. We calculated pathway scores for those using PROGENy, Reactome and Gene Ontology enrichment, SPIA, Pathifier, and PARADIGM. We used an ANOVA to calculate significant associations between the presence and absence of mutations and copy number alterations and the inferred pathway scores for our method (Fig. 3a) and others, both with and without regressing out cancer types (Supplemental Fig. 7).

**3.**
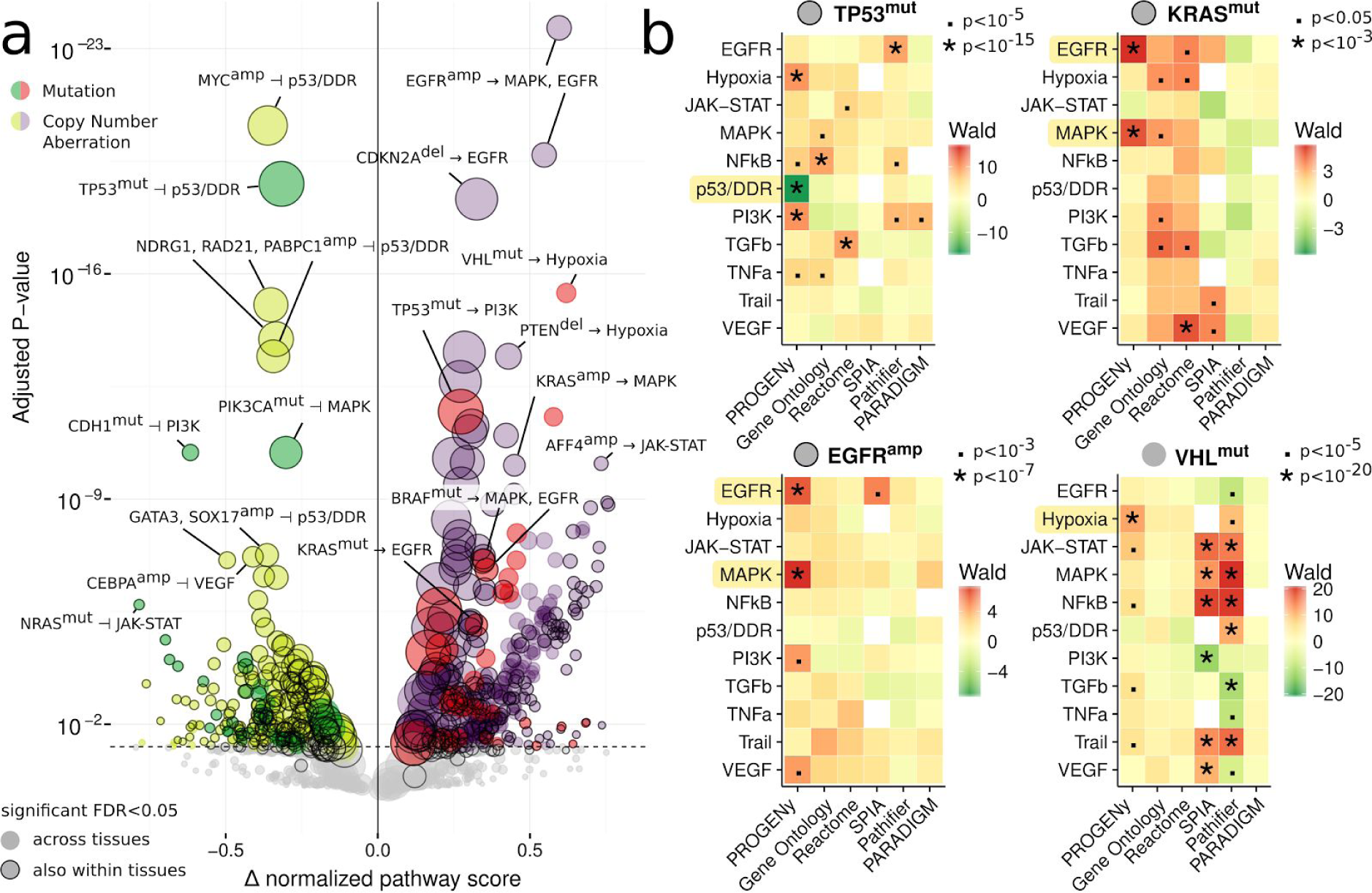
Ability of pathway methods to recover well-known mutations. A. Volcano plot of pan-cancer associations between driver mutations and copy number aberrations with differences in pathway score. Pathway scores calculated from basal gene expression in the TCGA for primary tumors. Size of points corresponds to occurrence of aberration. Type of aberration is indicated by superscript “mut” if mutated and “amp”/”del” if amplified or deleted, with colors as indicated. Effect sizes on the horizontal axis larger than zero indicate pathway activation, smaller than zero inferred inhibition. P-values on the vertical axis FDR-adjusted with a significance threshold of 5%. Associations shown without correcting for different cancer types. Associations with a black outer ring are also significant if corrected. B. Comparison of pathway scores (vertical axes) across different methods (horizontal axes) for *TP53* and *KRAS* mutations, *EGFR* amplifications and *VHL* mutations. Wald statistic shown as shades of green for downregulated and red for upregulated pathways. P-value labels shown as indicated. White squares where a pathway was not available for a method.

In terms of proliferative signaling, we find that PROGENy identifies *EGFR* amplifications to activate both the EGFR and MAPK pathways (FDR<10^−20^), and to a lesser extent PI3K, VEGF, and Hypoxia (FDR<10^−9^). *KRAS* mutations show an increase in inferred EGFR activity, and amplifications additionally for MAPK and PI3K (FDR<10^−5^). All other methods fail to detect a strong activation of the MAPK/EGFR pathways (Fig 3b.; top right and bottom left) given those alterations. We further find an increase in PI3K activity with *ERBB2* amplifications, but also a reduction in the Trail signature (FDR<0.05), suggesting a stronger relative impact on cell survival. *BRAF* mutations have a positive effect on EGFR and MAPK (FDR<10^−9^) but not PI3K (FDR>0.4).

For *TP53* mutations PROGENy finds a significant reduction in p53/DNA damage response activity (FDR<10^−18^) and activation of the pathways for MAPK, PI3K, and Hypoxia (FDR<10^−4^). This is in contrast to loss of *TP53*, where we only find a reduction in p53/DDRactivate both the EGFR and MAPK pathways (FDR<10^−3^) but no modification of any other pathway (FDR>0.15). The dual nature of *TP53* mutations and loss are in line with the recent discovery that *TP53* mutations can act in an oncogenic manner in addition to disrupting its tumor suppressor activity, which has been shown for individual cancer types (Olive et al. 2004; Zhang et al. 2013; Weissmueller et al. 2014; Zhu et al. 2015). In addition, this analysis suggests a link between *TP53* mutations and genes that are induced by activation of canonical oncogenic signaling such as MAPK and PI3K. Other methods (Fig 3b.; top left) are unable to recover the expected negative association between these alterations and p53/DDR activity. GO and Reactome showed a much weaker effect in the same direction, while Pathifier and SPIA showed an incorrect positive effect. These methods do, however, capture the activation of other oncogenic pathways, suggesting this effect is driven by expression changes that then leads to changes in activity.

PROGENy finds that *VHL* mutations (which have a high overlap with Kidney Renal Carcinoma, KIRC) are associated with an expected stronger induction of hypoxic genes (Maxwell et al. 1999) compared to other cancer types. It is the only one to recover hypoxia as the strongest link with *VHL* mutations, while the other methods primarily report expression changes in unrelated pathways (Fig 3b.; bottom right). More surprisingly, we find that presence of *PIK3CA* amplifications and *PTEN* deletions is also more connected to increasing the hypoxic response (FDR<10^−6^) compared to an effect on the PI3K-responsive genes (FDR between 10^−2^ and 10^−5^). A role of PI3K signaling in hypoxia has been shown before (Zhou et al. 2004; Yang et al. 2009; Kilic-Eren et al. 2013).

These highlights reflect the more general pattern PROGENy is able to correctly infer the impact of driver mutations that other pathway methods are not. The latter are only able to identify some cases where activity is mediated by changes in the expression level (Supplemental Table 7).

### Associations with drug response

The next question we tried to answer is how well PROGENy is able to explain drug sensitivity in cancer cell lines. We took as a measure of efficacy the IC_50_, i.e. the drug concentration that reduces viability of cancer cells by 50%, for 265 drugs and 805 cell lines from the GDSC project (Iorio et al. 2016) and performed an ANOVA between those and inferred pathway scores of PROGENy, Reactome, Gene Ontology, SPIA, Pathifier, and PARADIGM.

We found 199 significant associations (10% FDR in Fig. 4a and Supplemental Fig. 7) for PROGENy, dominated by sensitivity associations between MAPK/EGFR activity and drugs targeting MAPK pathway (Fig. 4b) that are consistent with oncogene addiction. In particular, this includes associations of the MAPK/EGFR pathways with different MEK inhibitors (Trametinib, RDEA119, CI-1040, etc.), a RAF inhibitor (AZ628) and a TAK1 inhibitor (7-Oxozeaenol). However, the strongest hit we obtained was the association between Nutlin-3a and p53-responsive genes. Nutlin-3a is an MDM2-inhibitor that in turn stabilizes p53, and it has also previously been shown that a mutation in *TP53* is strongly associated. with increased resistance to Nutlin-3a (Garnett et al. 2012). Hence, this is a wellunderstood mechanism of sensitivity (presence) or resistance (absence of p53 activity) to this drug that our method captures but none of the pathway expressionbased methods do.

**4.**
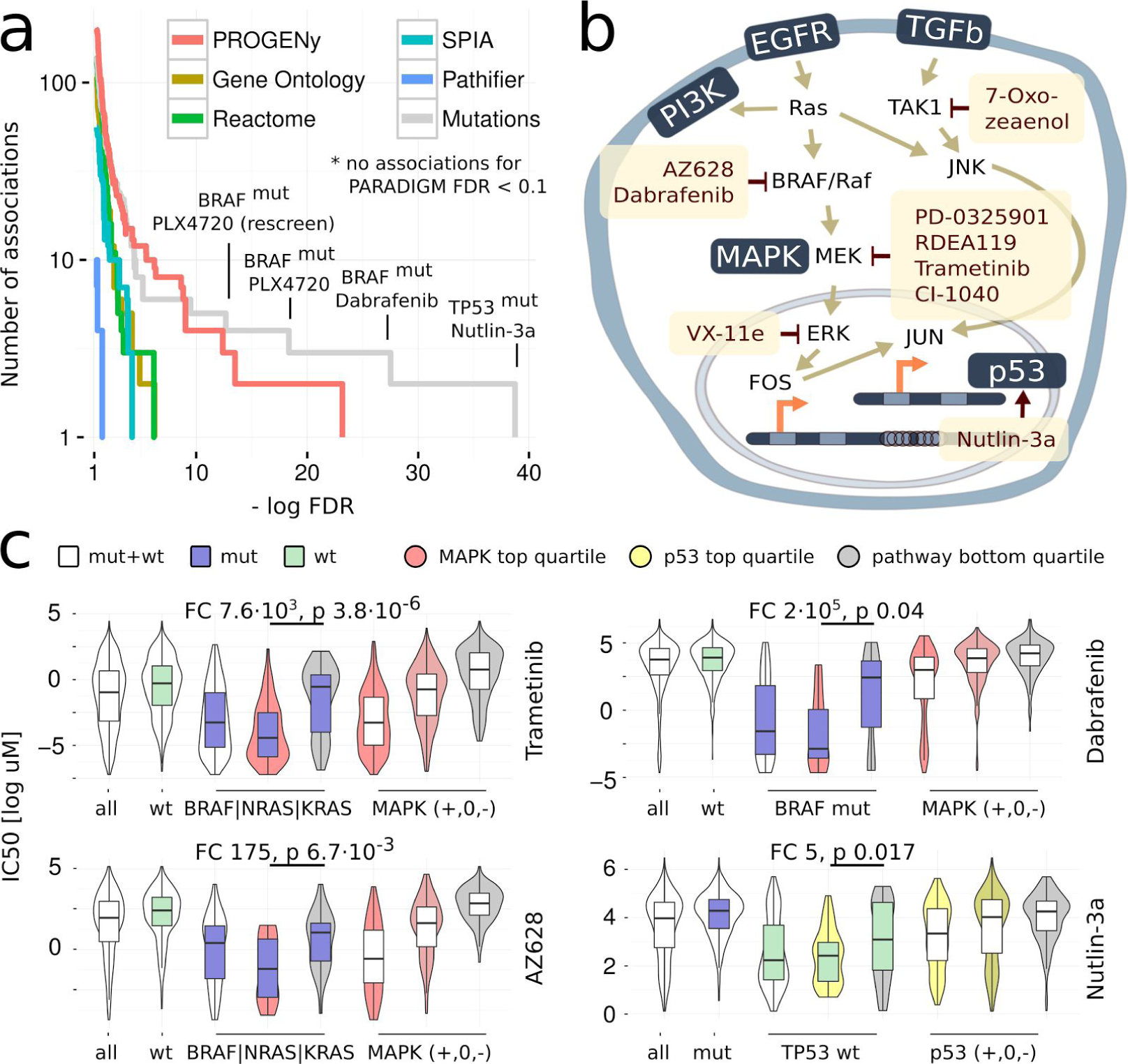
MAPK and p53 scores drive drug response across all cancer types. A. Comparison of the associations obtained by different pathway methods. Number of associations on the vertical, FDR on the horizontal axis. PROGENy yield more and stronger associations than all other pathway methods. Mutation associations are only stronger for TP53/Nutlin-3a and drugs that were specifically designed to bind to a mutated protein. PARADIGM not shown because no associations < 10% FDR. markers (green) and greater than zero resistance markers (red). P-values FDR-corrected. B. Pathway context of the strongest associations (Supplemental Fig. 8) between EGFR/MAPK pathways and their inhibitors obtained by PROGENy. C. Comparison of stratification by mutations and pathway scores. MAPK pathway ( *BRAF*,*NRAS*, or *KRAS*) mutations and Trametinib on top left panel, AZ628 bottom left, *BRAF* mutations and Dabrafenib top right, and p53 pathway/*TP53* mutations/Nutlin-3a bottom right. For each of the four cases, the leftmost violin plot shows the distribution of IC_50_s across all cell lines, followed by a stratification in wild-type (green) and mutant cell lines (blue box). The three rightmost violin plots show stratification of all the cell lines by the top, the two middle, and the bottom quartile of inferred pathway score (indicated by shade of color). The two remaining violin plots in the middle show mutated (*BRAF*, *KRAS*, or *NRAS*; blue color) or wild-type (*TP53*; green color) cell lines stratified by the top- and bottom quartiles of MAPK or p53 pathways scores (Mann-Whitney U test statistics as indicated).

Considering the overall number of associations, the other pathway methods provided a lower number across the range of significance (Fig. 4a and Supplemental Fig. 7). PROGENy even outperforms associations obtained with driver mutations at 10% FDR, as those only yield 136 associations. The latter only provide stronger associations for *TP53*, where the signature is a compound of p53 signaling and DNA damage response, and PLX4720/Dabrafenib, drugs that were specifically designed to target mutated *BRAF*. For 170 out of 265 drugs covered by significant associations with either PROGENy or driver mutations, PROGENy provided stronger associations for 85, with a significant enrichment in cytotoxic drugs compared to targeted drugs for mutations (Fisher’s exact test, p<0.002).

However, stratification using PROGENy and mutated driver genes is not mutually exclusive. Our pathway scores are able to further stratify the mutated and wild-type sub-populations into more and less sensitive cell lines (Fig. 4c and Supplemental Tables 8-9). This includes, but is not limited to, *BRAF*, *NRAS* or *KRAS* mutations using MAPK pathway activity and the MEK inhibitor Trametinib (Fig. 4c; top left) or RAF inhibitor AZ628 (Fig. 4c; bottom left), *BRAF* mutations with Dabrafenib (Fig. 4c; top right), and *TP53* mutations with p53/DDR and Nutlin-3a (Fig. 4c; bottom left). For MAPK- and *BRAF*-mutated cell lines, we find that cell lines with an active MAPK pathway according to the PROGENy are 175 (AZ628), 7596 (Trametinib), or 10^5^ fold (Dabrafenib) more sensitive than those where it is inactive. For Trametinib, cell lines with active MAPK but no mutation in *BRAF*, *KRAS*, or *NRAS* are six times more sensitive than cell lines that harbor a mutation in any of them but MAPK is inactive.

Taken together, these results show that PROGENy can be used to complement mutation-derived biomarkers by either refining them or providing an alternative where no such marker exists. Associations obtained with other methods do not show strong interactions between pathways and drugs that target their members.

### Implications for patient survival

The implications of inferred pathway activity compared to pathway expression is expected to be less clear for patient survival than for cell line drug response due the many more factors that affect the phenotype observed. Nonetheless, we were interested in how our inferred pathway activity compared to pathway expression methods in terms of overall patient survival.

Across all cancer types, PROGENy found a strong association with decreased survival for EGFR, MAPK, PI3K, and Hypoxia (Fig. 5a). Gene Ontology found much weaker associations for those pathways, and the other methods missed them almost entirely. In terms of Trail activity, PROGENy is the only method to find an increase in survival, while the other methods show either a decrease or no effect. For JAK-STAT, NFkB, p53, and VEGF there are no significant associations that are picked up by more than one method (FDR<0.05). In comparison, driver mutations did not provide any significant associations except for *TP53* (FDR<0.03 vs. FDR>0.2) with a weaker effect size compared to PROGENy.

**5.**
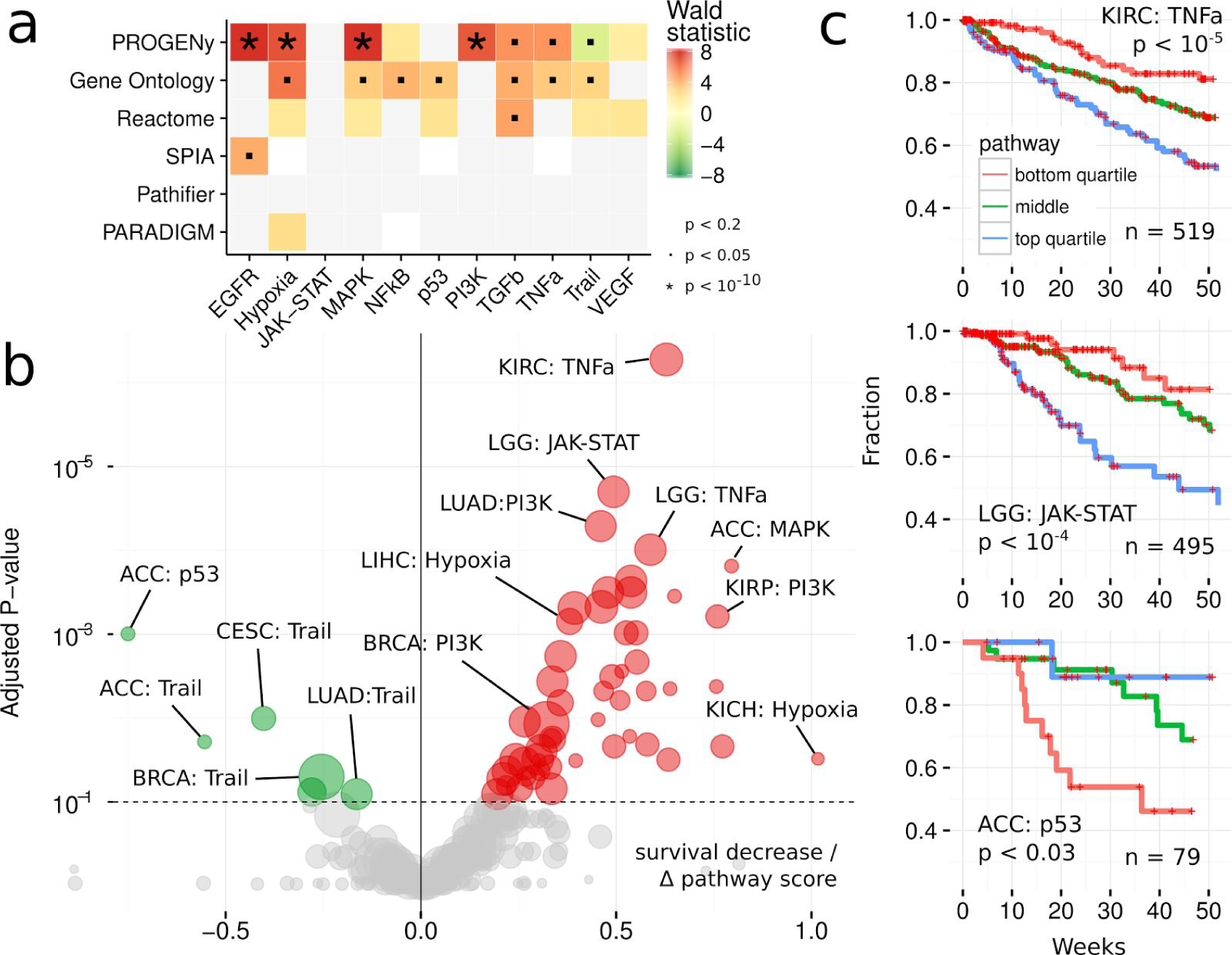
Response signatures outperform pathway methods for patient survival. A. Pan-cancer associations between pathway scores and patient survival. Pathways on the horizontal, different methods on the vertical axis. Associations of survival increase (green) and decrease. Significance labels as indicated. Shades correspond to effect size, p-values as indicated. B. Volcano plot of cancers-specific associations between patient survival and inferred pathway score using PROGENy. Effect size on the horizontal axis. Below zero indicates increased survival (green), above decreased survival (red). FDR-adjusted p-values on the vertical axis. Size of the dots corresponds to number of patients in each cohort. C. Kaplan-Meier curves of individual associations for kidney (KIRC), low-grade glioma (LGG) and adrenocortical carcinoma (ACC). Pathway scores are split in top- and bottom quartiles and center half. Lines show the fraction of patients (vertical axis) that are alive at a given time (horizontal axis) within one year. P-values for discretized scores.

For individual cancer types, PROGENy finds a similar separation between oncogenic and tumor-suppressor pathways (Fig. 5b) that other methods fail to provide (Supplemental Fig.9). Our associations are significant for more cancer types and more specific to individual pathways. We find cancer-specific associations of pathways with no effect in the pan-cancer setting. Adrenocortical Carcinoma (ACC) shows a significant survival increase with p53 activity (FDR<10^−3^), supported by the fact that it not harbor any previously reported gain-of-function *TP53* variants (Zhu et al. 2015). Kidney Renal Clear Cell Carcinoma (KIRC) and Low-Grade Glioma (LGG) show decreased survival with TNFa and JAK-STAT respectively, pathways where activating mutations are much less well established than for EGFR/MAPK. For these three associations, the top and bottom quartiles of PROGENy pathway activity were able to stratify patients in groups with over 25% difference in one year survival (Fig. 5c). Compared to mutations, PROGENy also provided stronger associations for cancer-specific survival (FDR 10^−7^ vs. 10^−3^).

## Discussion

The explanation of phenotypes in cancer, such as cell line drug response or patient survival, has largely been focussed on genomic alterations (mutations, copy number alterations, and structural variations). While this approach has generated many important insights into cancer biology, it does not directly make statements about the impact of those aberrations have on cellular processes and signal transduction in particular. Pathway methods, mostly used on gene expression, have so far largely fallen short on delivering actionable evidence. This can in part be due to lack of robustness, as suggested by the heterogeneity in responses of individual signatures (Fig. 2a), but arguably also by the fact that extracting features that reflect pathway activity from gene expression is not trivial. With proteomics lagging behind sequencing data for the foreseeable future, we have a need to address both the accurate inference of pathway activity from gene expression, as well as the issue of irreproducible gene signatures.

We developed PROGENy in order to overcome these limitations. PROGENy leverages a large compendium of pathway-responsive gene signatures derived from a wide range of different conditions in order to identify genes that are consistently deregulated. The result is a simple linear model that outperforms competing pathway methods that are orders of magnitude more complex and computationally expensive.

We found that despite the heterogeneity of individual gene expression experiments, PROGENy more closely corresponds to pathway perturbations than other methods. PROGENy can recover the impact of known driver mutations from basal gene expression, but also identify cases where a pathway is active without their presence. Pathway mapping only recovers known associations where this effect is mediated by expression changes, such as *TP53* oncogene activation or copy number aberrations.

In terms of drug sensitivity, we showed that PROGENy provides stronger associations than available pathway methods that also correspond better to known interactions. It can be used to refine mutation-derived biomarkers, as well as to provide novel markers with no associated mutation. Pathway expression is further removed and thus more likely to be a consequence rather than a cause of the drug sensitivity mediated by a signaling aberration. The fact that competing methods do not recover oncogene addiction patterns supports this claim.

For survival associations, only PROGENy finds the pathways that we would most expect to decrease patient survival by accelerating tumour growth (EGFR and MAPK) and promoting survival by apoptosis (Trail) to be associated with the respective outcome in both the pan-cancer as well as the tissue-specific cohorts. Other methods fail to separate those, only obtain significant associations for a very limited number of cancer types, and show high correlation between pathways.

Overall, our results suggest that PROGENy provides a better measure of pathway activity than other pathway methods, irrespective of whether the latter was derived from gene sets or directed paths. We have shown that PROGENy is able to refine our understanding of the impact of mutations, as well as their utility for cell line drug response and patient survival. It provides a strong evidence that in order to infer pathway activity, a downstream readout should be used instead of mapping transcript expression levels to signaling molecules.

## Methods

### Data from The Cancer Genome Atlas (TCGA)

To obtain TCGA data, we used the Firehose tool from the BROAD institute ( http://gdac.broadinstitute.org/), release 2016_01_28.

For gene expression, we used all data labelled “Level 3 RNA-seq v2”. We extracted the raw counts from the text files for each gene, discarded those that did not have a valid HGNC symbol, and averaged expression levels where more than one row corresponded to a given gene. We then performed a voom transformation (limma package, BioConductor) for each TCGA study separately, to be able to use linear modeling techniques with the count-based RNA-seq data. The data used corresponds to 34 cancer types and a total of 9737 tumor and 641 matched normal samples.

From clinical data, we extracted the vital status and used known survival time or known time of last follow-up as the survival time for the downstream analyses. We converted the time in days to months by dividing by 30.4. The overlap of TCGA data where we could obtain both mRNA expression levels as well as survival times is 10544 distributed across 33 cancer types. For comparing different pathway methods, we only used cancer types with tissue-matched controls, leaving 5927 samples in 13 cancer types.

### Data from the Genomics of Drug Sensitivity in Cancer (GDSC) project

We used version 17a of the GDSC data (Iorio et al. 2016), comprised of molecular data for 1,001 cell lines and 265 anticancer drugs, specifically microarray gene expression data (ArrayExpress accession E-MTAB-3610) and the IC_50_ values for each drug-cell line combination. For computing pan-cancer associations, we used the subset with TCGA-like cancer type label, leaving 768 cell lines.

### Curation of Perturbation-Response Experiments

Our method is dependent on a sufficiently large number publicly available perturbation experiments that activate or inhibit one of the pathways we were looking at. The following conditions needed to be met in order for us to consider an experiment: (1) the compound or factor used for perturbation was one of our curated list of pathway-perturbing agents (Supplemental Note 2); (2) the perturbation lasted for less than 24 hours to capture genes that belong to the primary response; (3) there was raw data available for at least two control arrays and one perturbed array; (4) it was a single-channel array; (5) we could process the arrays using available BioConductor packages; (6) the array was not custom-made so we could use standard annotations.

We curated a list of known pathway activators and inhibitors for 11 pathways, where the interaction between each compound and pathway is well established in literature. We then used those as query terms for public perturbation experiments in the ArrayExpress database (Parkinson et al. 2007) and included a total of 223 submissions and 573 experiments in our data set, where each experiment is a distinct comparison between basal and perturbed arrays. If there were multiple time points, different cells, different concentrations, or different perturbing agents within a single database submission, they were considered as different experiments.

### Microarray Processing

Started from the curated list of perturbation-induced gene expression experiments, we included all single-channel microarrays with at least duplicates in the basal condition with raw data available that could be processed by either the limma (Smyth 2005), oligo(Carvalho and Irizarry 2010), or affy (Gautier et al. 2004) BioConductor packages and for which there was a respective annotation package available. Multiple concentrations or time points in a series of arrays were considered as individual experiments.

We first calculated a probe-level for 573 full series of arrays, where we performed quality control of the raw data using RLE and NUSE cutoffs under 0.1 and kept all arrays below that threshold. If after filtering less than two basal condition arrays remained, the whole experiment was discarded. For the remaining 568 series we normalized using the RMA algorithm and mapped the probe identifiers to HGNC symbols.

### Building a Linear Model of Pathway-Response Genes

For each HGNC symbol, we calculated a model based on mean and standard deviation of the gene expression level, and computed the z-score as average number of standard deviations that the expression level in the perturbed array was shifted from the basal arrays. We then performed LOESS smoothing for all z-scores in a given experiment using our null model, as described previously (Parikh et al. 2010).

From the z-scores of all experiments and all pathways, we performed a multiple linear regression with the pathway as input and the z-scores as response variable for each gene separately:

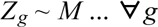

Where *Z_g_* is the z-score for a given gene *g* across all input experiments (as a column vector of experiments). *M* is a coefficients matrix (rows are experiments, columns pathways, Fig.1b) that has the coefficient 1 if the the experiment had a pathway activated, −1 if inhibited, and 0 otherwise. For instance, the Hypoxia pathway had experiments with low oxygen conditions set as 1, HIF1A knockdown as −1, and all other experiments as 0. The same is true for EGFR and EGF treatment vs. EGFR inhibitors respectively, with the additional coefficients of MAPK and PI3K pathways set to 1 because of known cross-talk (for a full structure of the cross-talk modeled, see Fig. 1c). As these are fold changes, we do not allow an intercept.

From the result of the linear model, we selected the top 100 genes per pathway according to their p-value and took their estimate (the fitted z-scores) as coefficient. We set all other gene coefficients to 0, so this yielded a matrix with HGNC symbols in rows and pathways in columns, where each pathway had 100 non-zero gene coefficients (Supplemental Table 1).

### PROGENy scores

Each column in the matrix of perturbation-response genes corresponds to a plane in gene expression space, in which each cell line or tumor sample is located. If you follow its normal vector from the origin, the distance it spans corresponds to the pathway score *P* each sample is assigned (matrix of samples in rows, pathways in columns). In practice, this is achieved by a simple matrix multiplication between the gene expression matrix (samples in rows, genes in columns, values are expression levels) and the model matrix (genes in rows, pathways in columns, values are our top 100 coefficients):

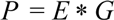

We then scaled each pathway or gene set score to have a mean of zero and standard deviation of one, in order to factor out the difference in strength of gene expression signatures and thus be able to compare the relative scores across pathways and samples at the same time.

### Pathway and Gene Ontology scores

We matched our defined set of pathways to the publicly available pathway databases Reactome (Croft et al. 2011)and KEGG (Kanehisa and Goto 2000), and Gene Ontology(GO) (Gene Ontology Consortium 2004) categories (Supplemental Tables 2-3), to obtain a uniform set across pathway resources that makes them comparable. We calculated pathway scores as Gene Set Variation Analysis (GSVA) scores that are able to assign a score to each individual sample (unlike GSEA that compares groups).

### SPIA scores

Signaling Pathway Impact Analysis (SPIA) (Tarca et al. 2008) is a method that utilizes the directionality and signs in a KEGG pathway graph to determine if in a given pathway structure the available species are more or less available to transduce a signal. As the species considered for a pathway are usually mRNAs of genes, this method infers signaling activity by the proxy of gene expression. In order to do this, SPIA scores require the comparison with a normal condition in order to compute both their scores and their significance.

We used the SPIA Bioconductor package (Tarca et al. 2008) in our analyses, focussing on a subset of pathways (Supplemental Table 4). We calculated our scores either for each cell line compared to the rest of a given tissue where no normals are available (i.e. for the GDSC and drug response data) or compared to the tissue-matched normals (for the TCGA data used in driver and survival associations).

### Pathifier scores

As Pathifier (Drier et al. 2013) requires the comparison with a baseline condition in order to compute scores, we computed the GDSC/TCGA scores as with SPIA. As gene sets, we selected Reactome pathways that corresponded to our set of pathways (Supplemental Table 3), where Pathifier calculated the “signal flow” from the baseline and compared it to each sample.

### PARADIGM scores

We used the PARADIGM software from the public software repository (https://github.com/sbenz/Paradigm) and a model of the cell signaling network (Cancer Genome Atlas Network 2012) from the corresponding TCGA publication (https://tcga-data.nci.nih.gov/docs/publications/coadread_2012/). We normalized our gene expression data from both GDSC and TCGA using ranks to assign equally spaced values between 0 and 1 for each sample within a given tissue. We then ran PARADIGM inference using the same options as in the above publication for each sample separately. We used nodes in the network representing pathway activity to our set of pathways (Supplemental Table 5) to obtain pathway scores that are comparable to the other methods, averaging scores where there were more than one for a given sample and node.

### Associations with known driver mutations and CNAs

For comparing the impact of mutations across different pathway methods, we used TCGA cohorts where tissue-matched controls were available, leaving 6549 samples across 13 cancer types. For mutated genes, we considered all genes that had a change of coding sequence (SNP, small indels in MAF files) as mutated and all others as not mutated. For copy number alterations (CNAs), we used the thresholded GISTIC (Beroukhim et al. 2007) scores, where we considered homozygous deletions (-2) and strong amplifications (2) as altered, no change (0) as basal and discarded intermediate values (-1, 1) from our analysis. We focussed our analysis of the mutations and copy number alterations on the subset of 464 driver genes that were also used in the GDSC. We used the sets of mutations and CNAs to compute the linear associations between samples for all different methods we looked at. We did not regress out the cancer type in order to keep associations where mutations/CNAs are highly correlated with it, but highlighted all associations that passed the significance threshold of FDR<5% (for each pathway method individually) after such a correction.

### Drug associations using GDSC cell lines

We performed drug association using an ANOVA between 265 drug IC_50_s and 11 inferred pathway scores conditioned on MSI status, doing a total of 2915 comparisons for which we correct the p-values using the false discovery rate. For pan-cancer associations, we used the cancer type as a covariate in order to discard the effect that different tissues have on the observed drug response. While this will also remove genuine differences in pathway activation between different cancer types, we would not be able to distinguish between those and other confounders that impact the sensitivity of a certain cell line from a given tissue to a drug. Our pan-cancer association are thus the same of intra-tissue differences in drug response explained by inferred (our method, GO, or Reactome) pathway scores.

We also selected two of our strongest associations to investigate whether they provide additional information of what is known by mutation data. For two MEK inhibitors, we show the difference between wild-type and mutant MAPK pathway (defined as a mutation in either *NRAS*, *KRAS*, or *BRAF*) with a discretized pathway score (upper and lower quartile vs. the rest), as well as the combination between the upper quartile of tissue-specific pathway scores and presence of a MAPK mutation.

### Survival associations using TCGA data

Starting from the pathway scores derived with GO/Reactome GSEA, SPIA, Pathifier, PARADIGM, and our method on the TCGA data as described above, we used Cox Proportional Hazard model (R package survival) to calculate survival associations for pan-cancer and each tissue-specific cohort. For the pan-cancer cohort, we regressed out the effect of the study and age of the patient, and fitted the more for each pathway and method used. For the tissue-specific cohorts, we regressed out the age of the patients. We adjusted the p-values using the FDR method for each method and for each method and study separately. We selected a significance threshold of 5 and 10% for the pan-cancer and cancer-specific associations for which we show a matrix plot and a volcano plot of associations, respectively.

In order to get distinct classes needed for interpretable Kaplan-Meier survival curves (Fig. 4c), we split all obtained pathway scores in upper, the two middle, and lower quartile and respectively to show for the three examples of associations found.

## Code availability

PROGENy [will be] available as an R package on Bioconductor. The code used for the analysis in this paper is available at https://github.com/saezlab/footprints

## Acknowledgements

### Acknowledgements

MS is funded by a MRC Case fellowship awarded to JSR and Joanna Betts (GSK). NB acknowledges funding by BMBF (OncoPath). MJG is supported with funding from the Wellcome Trust (102696), Stand Up To Cancer (SU2C-AACR-DT1213), The Dutch Cancer Society (H1/2014-6919) and Cancer Research UK (C44943/A22536). We thank Francesco Iorio, Florian Markowetz and Alvis Brazma for useful discussions.

## Author contributions

MS designed research, performed all analyses, and wrote the manuscript. BK, MK, NB and MJG supported result interpretation. JSR supervised the project and contributed to writing the manuscript.

